# Energetic benefits of social information for movement in patchy landscapes

**DOI:** 10.64898/2025.12.18.695131

**Authors:** Eleonora Gatti, Andreagiovanni Reina, Hannah J. Williams

## Abstract

Movement is costly, and animals are under strong selective pressure to move efficiently, yet, in patchy, dynamic landscapes, decision-making is inherently uncertain. We quantify the energetic savings achieved by using up-to-date information presented within social cues for reducing movement costs. We use an agent-based model, founded on realistic aeronautical rules and parametrised on the Andean condor (*Vultur gryphus*), to study movement in patchy landscapes. By explicitly considering altitude, flight results in a sequence of soaring and gliding in the 3D space. We investigate how the cost of movement to an overall goal varies when birds use social information from others that are either fixed in space or moving collectively to the common goal, and under different risk-taking speed strategies, from slow and cautious to fast and risky. The value of social information is operationalised as energetic savings in units of basal metabolic rate. Under low predictability, agents with intermediate risk and high social-information use exhibit lowest movement costs, with up to 41% energy savings over asocial movement. By extending classical aeronautical theory to social and variable environments we demonstrate the adaptive value of social information for efficient movement in patchy, unpredictable landscapes.

## 1. Introduction

Animals need to move to access resources required for life, thus, movement strategies are under strong selective pressure to be efficient [1], i.e. to reach intended destinations with minimal energy consumption. This need for efficiency has lead to species evolving mechanisms by which they can exploit energy available in the environment for cheap locomotion [2], which in some cases may be extreme. Soaring species, for example have evolved low wing loading (large wings relative to body mass) morphologies to maximise soaring potential for height gain using air currents, allowing them to glide and travel efficiently with minimal need to resort to expensive flapping flight. The harnessing of environmental energy significantly reduces the cost of transport—a metric that quantifies the amount of energy needed to move a given mass over a given distance—and has been adopted by various species across media via morphological adaptation. However, optimal movement decisions for energy-efficient paths in moving media are complex given uncertainty in the spatio-temporal dynamics of energy availability.

Adaptive behaviour depends on an individual’s capacity to use the available information to mitigate uncertainty; in the search for resources, the behaviour of conspecifics can provide the most relevant up-to-date information on resource availability [3]. Whereas social information is known to increase the efficiency of foraging [4, 5, 6, 7]—e.g. for vultures to locate carcasses [8], bats to find insect swarms [9], or whales to track ephemeral krill aggregations [10]—most studies have considered its role only in the context of food resource detection. Much like the value of information in locating ephemeral food patches in optimal foraging theory, information is key in movement decisions for reduced energetic expenditure [11]. Social information can be used to identify energy-efficient migratory paths via collective sensing or social learning [12, 13], and a growing body of work has also shown how the movement of individuals influences the behaviour of others, often described broadly as the “social environment” [14, 15, 16]. Thus, energetically expensive or short-sighted movement decisions can be avoided by considering up-to-date information about environmental energy resources (e.g. media flow) in the movements of others [11]. By reducing uncertainty about resource distribution and availability, social information improves decision-making and movement efficiency, i.e. lower energy consumption. Thus, social information is also a resource with an energetic value.

Minimising energy consumption is fundamental to ecological success [11, 17] and to determine the significance of social information to reduce uncertainty, we should quantify its energetic value. Recognising social information’s energetic value is vital, not only because it differs fundamentally from direct energetic effects of fluid dynamics when moving with conspecifics in specific formation or phase alignment [18, 19], but also for understanding the selective pressures that promote sociality; in the context of gaining access to foraging resources and in reducing the energetic costs of movement. Assessing how social information shapes movement decisions and characterising their energetic consequences can enable a quantitative integration of the social and the energy landscapes for movement analyses.

Here, we build a computational model to quantify the energetic value of social information for movement in the case of soaring-gliding flight of the heaviest of soaring birds, the Andean condor (*Vultur gryphus*). Efficient soaring flight relies on accurate movement decisions to move between sources of lift in a complex and uncertain environment, where balancing exploitation and exploration of energy patches is essential for cross-country flight. In the dynamic aerial environment with invisible media flows, detecting patches of lift is challenging and relies heavily on visual-based sensing of underlying cues related to probability of lift formation [20]. Landscape features and weather conditions may offer indirect cues, but these alone are often insufficient to predict where and when thermals will be active at any given moment. The presence and movement of other soaring individuals are a reliable source of up-to-date information, reducing uncertainty on precise resource location and characteristics [21, 22, 23], making this a model system to quantify the energetic savings that can be made by using social information. Additionally, the consequence of an incorrect movement decision can be extreme. Poor decisions can result in costly outcomes (up to 20 times basal metabolic rate [24]), where the bird must resort to flapping to reach active patches of lift [25]. Therefore, the energetic benefits of using social information can be significant, creating strong selective pressures for efficient movement strategies [26, 27].

Accurate decision-making in soaring flight manifests in the selection of gliding speed, where flying faster enables quicker access to resources but comes at the cost of increased altitude loss and, therefore, higher risk of having to resort to expensive flapping flight to avoid landing. According to aeronautical theories, a soaring individual’s gliding speed and sink rate (rate of altitude loss per unit time) are linked by a theoretical curve, the glide polar [28, 29]; where increased airspeed in the glide results in a greater sink rate. That is, individuals must choose between opting for a slow and cautious glide that conserves altitude, i.e. energy reserves, or a fast and risky glide that increases cross-country speed (average ground speed between two points) [30]. Informed movement decisions should better balance the trade-off between time efficiency and altitude loss, given the environmental predictability. Moreover, birds exploiting thermals discovered by others leads to collective movement, where networks of individuals share information and collectively sense the environment to track ephemeral invisible energy resources.

To quantify the energetic value of social information on movement performance in our soaring system, we created an agent-based model that simulates soaring individuals navigating a theoretical energy landscape, offering a flexible, bottom-up approach to tackle ecological problems [31, 32, 33]. In our model, the environment comprises a distribution of resource patches that allow birds to gain altitude and move across the landscape. Movement is defined as a series of glide-soar sequences between updraughts (patches of lift), each with an energetic cost in terms of time and flapping required to avoid landing, determining overall flight performance. The agent is characterised by two key parameters: (1) sociality *s*, a value that represents the extent to which movement decisions are influenced by social information, and (2) risk *r*, representing the tendency to expend energy reserves (height) by selecting higher glide speeds on the polar glide curve. In soaring flight, social information comes from the presence of conspecifics in active updraughts, which signals the availability of energy resources. Agents are pulled towards the same spatially well-defined goal, which can represent, for example, a known food location or an established roosting site. With this model, we measure performance in terms of energetic costs—specifically the time spent gliding and flapping to reach the goal—across different simulated environments where the predictability and availability of energy resources varied. We simulate two scenarios: (1) a single focal individual moving in a fixed social environment, and (2) a collective of homogeneous individuals. This design allows us to separate the effects of uniformly distributed social information from those arising through collective movement, which generates dynamic and spatially heterogeneous social cues.

In general, our model allows for quantifying the energetic savings that result from increased sociality, thus assigning an energetic value to social information. We hypothesise that greater social information use will exhibit higher performance, especially in challenging environments where resources are scarce. In contrast, higher risk-taking may lead to increased chances of failure regardless of environmental conditions, as higher glide speeds are associated with greater sink rates, increasing the likelihood of reaching the ground before encountering a suitable thermal [30]. For the collective simulations, we expect an explore-exploit trade-off between movement towards the overall goal and and attraction to socially indicated local patches of lift [34], such that collective movement may not always yield energetic savings.

## 2. Methods

### 2.1 Model description

We built an agent-based model simulating the flight of the Andean condor (*Vultur gryphus*), the heaviest obligate soaring species, where flight performance and limits were based on the species’ morphology according to [28]. The model was implemented in Julia v”1.10.2”, using the package Agents.jl [35]. A list of the model parameters, their meaning and units are available in Table 1. The model can simulate either a single or a group of soaring individuals moving through a patchy energy landscape of lift (hereafter thermals).

**Table 1:** List of model parameters with their description and values or variation range.

| Name | Notation | Description | Value/Range (units) |
| --- | --- | --- | --- |
| Sociality | $s$ | Reliance on social information | [0, 1] |
| Risk | $r$ | Tendency to choose higher glide speeds | [0, 1] |
| Predictability | $p$ | Probability that a thermal is active | [0, 1] |
| Availability of social information | $a_{si}$ | Probability that an active thermal is occupied | [0, 1] |
| Number of thermals | $N_{th}$ | Number of thermals in the environment | [1111, 3265, 40000] |
| Best glide speed | $v_{bg}$ | Glide speed for maximum distance | 14.8 m/s |
| Best glide ratio | $gr_{max}$ | Glide ratio associated with $v_{bg}$ | 16.2 |
| Maximum glide speed | $v_{max}$ | Upper limit of glide speed | 25.0 m/s |
| Fixed sink rate | $sink$ | Rate of altitude loss when not gliding | 1.0 m/s |
| Dimensions | $dims$ | Dimensions of the arena | 100 km × 100 km |
| Goal position | $goal$ | Coordinates of the goal | (99, 99) |
| Thermal quality | $q$ | Quality of a thermal as vertical wind speed | [1, 4] m/s |
| Variance of directed distribution | $\sigma_{dir}$ | Standard deviation of directed probability distribution | 1500 |
| Weight factor | $K$ | Weight factor used to combine Gaussian and directed probability distributions | 0.5 |
| Maximum height | $h_{max}$ | Maximum height that the bird can possibly reach | 1500 m |

#### 2.1.1 Environment

Movement took place in a continuous two-dimensional space of 100 *×* 100 km^2^, containing *N*_th_ thermals (Fig. 1A). Each thermal had a fixed position and a quality value *q*, determining the bird’s altitude gain when exploiting it (i.e. vertical velocity). Thermals could be active (providing lift) with a probability represented by a parameter predictability *p* ∈ [0, 1], or inactive, not providing lift, despite being in a location where uplift was likely to be present. In the focal-individual scenario, with social information available with probability *a*_si_ ∈ [0, 1], each active thermal was additionally assigned an occupancy state, representing the presence of conspecifics. In this setup, the individual navigates an environment with static social information, where occupancy of thermals does not change throughout the simulation. In the collective simulation scenario, occupancy of a given thermal was updated dynamically based on the presence of birds soaring within the thermal.

**Figure 1:**
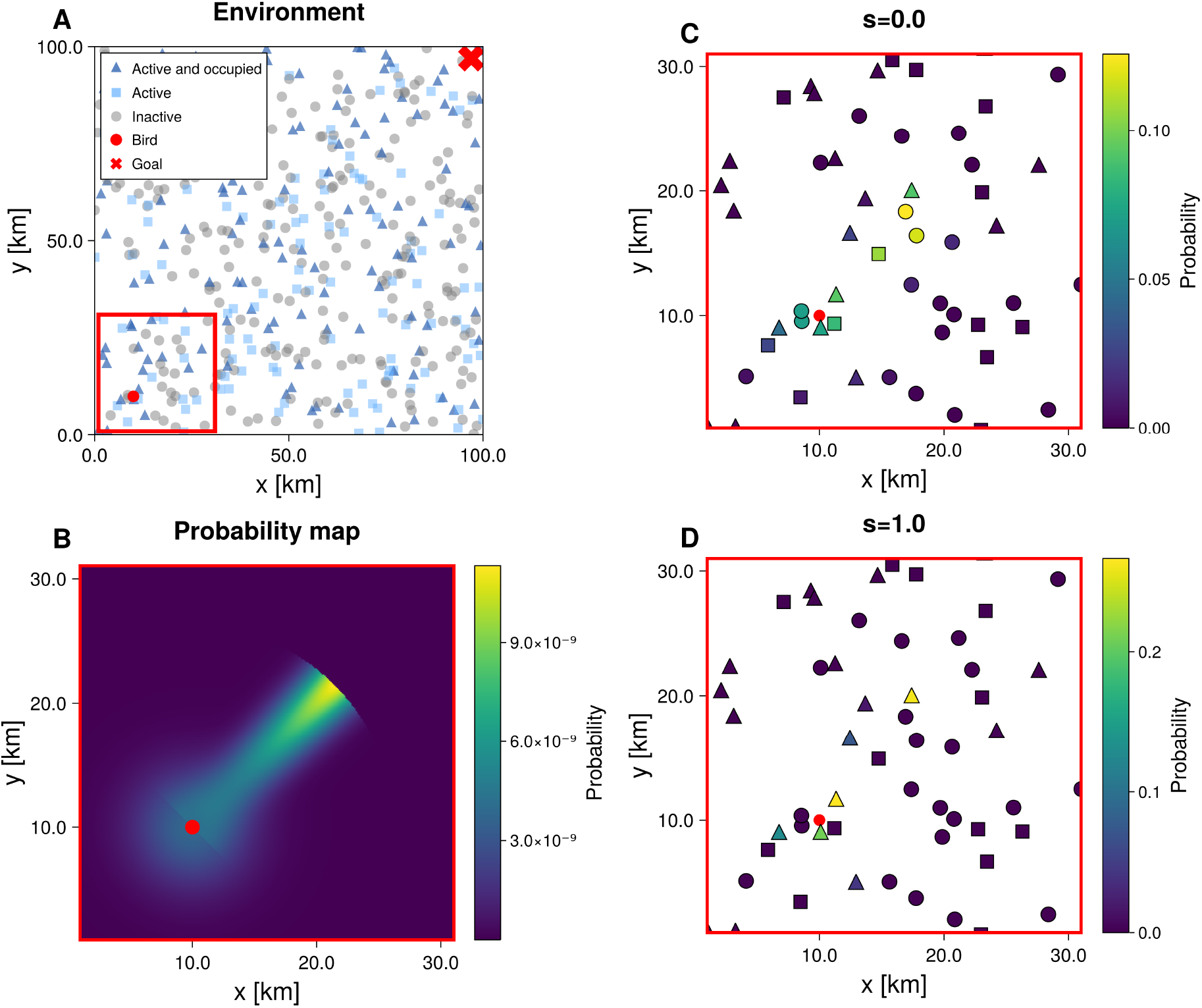
Workflow of the model to evaluate the probability of moving to the next location at each time step (example for simulation of an individual in a static social environment). The environment (panel A) is represented as a set of thermals that can be active (light blue squares), active and occupied (dark blue triangles), or inactive (grey circles); the red dot represents the position of the bird, and the red cross the location of the goal. The probability map (panel B) around the bird’s position is used to evaluate the probability assigned to each thermal (Eq. 1) and is then adjusted by the parameter *s* for calculating final probabilities (Eq. 4). We show examples of the extreme cases of sociality: when *s* = 0 (panel C), all reachable thermals contribute equally regardless of their occupation state; when *s* = 1 (panel D), only occupied thermals are considered. Based on these final probabilities, the next location is selected probabilistically.

#### 2.1.2 Bird movement

Each bird was defined by its position, altitude (*h*), and movement state. Depending on its movement state, the simulator updates the bird’s position and altitude every 10 seconds. The bird can be in either of three movement states: soaring, gliding, or searching.

- **Soaring**: the bird gains altitude within a thermal proportional to its quality. With a probability that increases with the thermal quality, the bird decides to continue soaring for the next 10 seconds. For low thermal quality, there is a higher probability that the bird will begin searching, select another thermal, and leave.
- **Gliding**: the bird moves towards the selected thermal while losing altitude at a rate determined by the glide polar.
- **Searching**: the bird searches and selects the next thermal within its effective radius *R* = *h · gr*_max_, where *gr*_max_ was the glide ratio at *v*_bg_ (the best glide speed) and hence, the maximum horizontal distance that could possibly be reached in a single glide. This effective radius does not represent the bird’s full perception range, which is larger and influenced by multiple factors [33], but instead defines the subset of locations that are both detectable and reachable given its current height and glide performance. The probability of selecting a thermal is influenced by personal information of the spatial distribution of thermals and by social information on current thermal activity (see *Sociality* below).

While in an active thermal *i*, the bird soars and gains *q·* 10 metres per time step. At each step (Δ*t* = 10 s), it either continues soaring with a probability that increases with thermal quality *q* (equivalently, *P*_stay_ = 1 − 1*/*5*q*), or it performs a one-step search. In the one-step search, the bird samples a destination thermal; if it selects the current thermal (th = *i*), it returns to soaring, otherwise it switches to gliding toward the selected thermal. When the bird has reached the maximum height, i.e. *h* ≥ *h*_max_, it cannot select the current thermal and is forced to leave. Upon arrival, if the target thermal is active, the bird resumes soaring; if inactive, it performs a one-step search.

#### 2.1.3 Decision-making

At each one–step search, a bird makes two probabilistic decisions: (i) which thermal to fly to, and (ii) the gliding speed to use. These decisions are modulated by two behavioural parameters:

- **Sociality** *s*: increases the probability of selecting thermals that are occupied, thus regulating reliance on social information.
- **Risk** *r*: shapes the distribution of gliding speeds, with higher values of risk corresponding to faster flight speeds along the glide polar but also increased sink rate and thus altitude loss.

##### Sociality and thermal selection

The probability of selecting a thermal at position **x** from the current position **c** is given by the weighted sum of two spatial components:

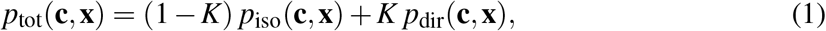

illustrated in Fig. 1B. The first component *p*_iso_(**c, x**) is an isotropic Gaussian kernel centred on **c**, representing undirected local exploration:

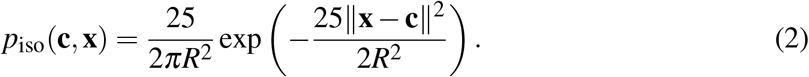

The second component *p*_dir_(**c, x**) is a directed kernel that increases the probability of selecting a thermal aligned with the vector from **c** to the goal, while down-weighting those thermals offset from this direction:

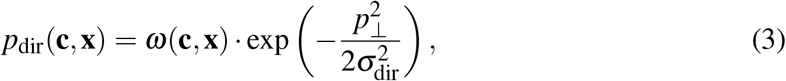

where *p*_⊥_ is the orthogonal projection of the vector from the current position **c** to a thermal in position **x** with respect to the goal direction, *σ*_dir_ is the spread of the directional bias, and *ω* is a distance-based weighting function proportional to ||**x** − **c**|| favouring thermals closer to the goal. The weight *K* in Eq. (1) sets the relative contribution of these two components. *K* was set to 0.5 as a balance between directed movement toward the goal and local exploration, which ensured both a reasonable success rate (see below) and computational efficiency. Candidate thermals are restricted to those within the effective radius *R*. Sociality modifies the resulting distribution as the probability of selecting an unoccupied thermal is reduced by a factor of 1 −*s*:

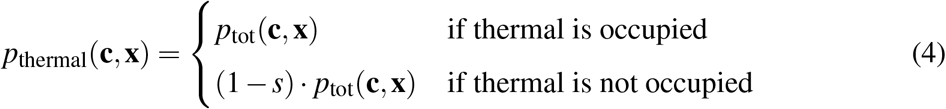

Thus, when *s* = 0 (no social influence), the weighting is neutral (Fig. 1C), while when *s* = 1 (fully social) unoccupied thermals are removed from the set of possibilities, leaving only occupied thermals as options (Fig. 1D). The new adjusted probabilities are then normalised. The thermal currently in use is always included among the available options if active and if *h < h*_max_. Finally, the probability given by Eq. (4) is sampled to select the next target thermal.

##### Risk and gliding speed

Once a thermal is chosen, the feasible range of gliding airspeeds is bounded by the best glide speed *v*_bg_ (that maximises both the horizontal distance per unit of height and the arrival altitude at the thermal) and the maximum speed *v*_max_ that allows reaching the thermal in the shortest time at positive altitude. Allowing glide speeds slower than *v*_bg_ would result in both greater height loss for a given horizontal displacement and longer glide time, which would be suboptimal in this context. The actual glide speed is sampled from a Beta distribution scaled to [*v*_bg_, *v*_max_], with shape parameters determined by *r* (Fig. 2). Low values of *r* yield distributions skewed toward *v*_bg_ (conservative choices), whereas high *r* increases the probability of selecting speeds closer to *v*_max_ (risky choices are associated with greater altitude loss). The glide polar (Fig. 2A) determines the sink rate *v*_z_ and glide ratio *gr* at the selected speed, which update altitude and effective radius.

**Figure 2:**
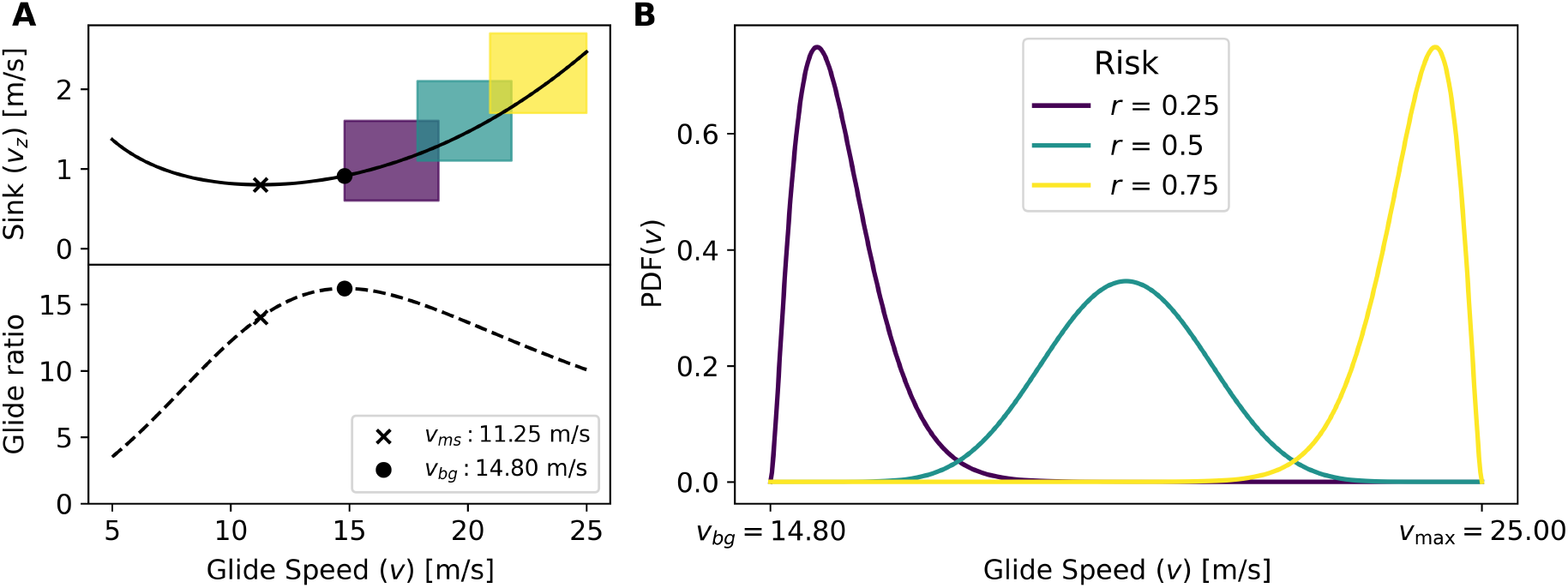
Glide polar curve parametrised with Andean condor (*Vultur gryphus*) parameter values (Table 2) (panel A). The coloured boxes represent the strategies selected based on different risk levels. The minimum sink speed *v*_ms_ minimises vertical sink rate per time unit, while the best glide speed *v*_bg_ minimises altitude loss per unit distance. The gliding speeds are selected according to the probability density function of the speeds (panel B). Lower values of *r* correspond to lower values of glide speed which are associated with a lower sink rate, thus resulting in a slower but safer movement. In contrast, higher values of *r* correspond to higher values of glide speed, resulting in quicker and more risky movements.

### 2.2 Experimental setup

We adopted a two-step approach. First, we performed a sensitivity analysis across the full parameter space to characterise model behaviour and guide the selection of fixed values for *a*_si_ and *N*_th_. Fixing these parameters allowed us to control the amount of social information via the predictability parameter. In a second step, we focused on testing our main hypotheses by varying the key parameters—sociality (*s*) and risk (*r*)—under different levels of environmental predictability (*p*).

**Table 2:**
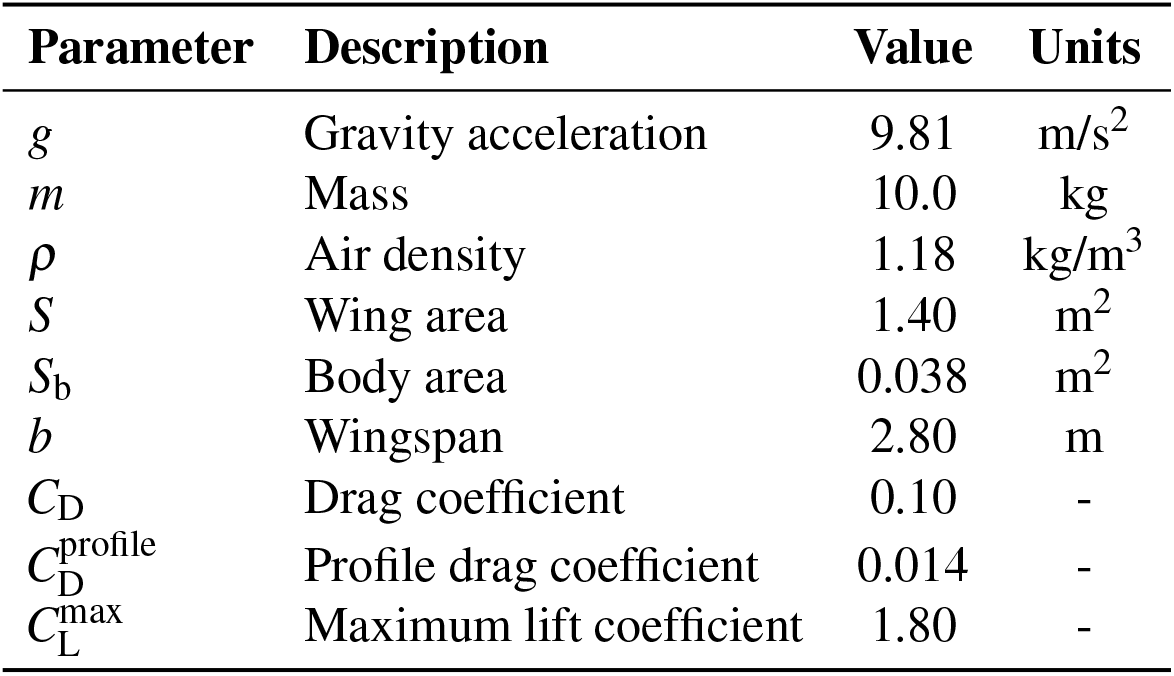
Values of parameters used to calculate the glide polar curve.

#### Sensitivity analysis

We initially explored five parameters in a sensitivity analysis: *s, r, p, a*_si_, and *N*_th_. We varied the parameters *s, r, p*, and *a*_si_ between 0 and 1, where individual state parameters *s* and *r* were scanned at intervals of 0.2 and the environmental parameters of *p* and *a*_si_, were scanned more sparsely, initially (in a subsequent analysis, *p* was explored in greater detail). For the parameter *N*_th_, representing the number of thermals in the environment, we analysed three realistic scenarios corresponding to increasing average distances between thermals (i.e. decreasing thermal density). Assuming that the average distance 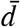 between thermals in an area with density of thermals *ρ* can be approximated by:

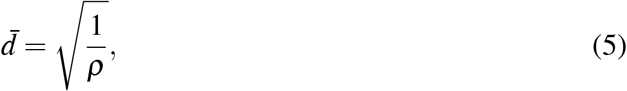

and given that 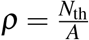, we derived

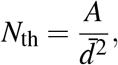

where *A* is the area of the simulated surface, 100 *×* 100 km^2^ (*A* = 10^10^ m^2^). The corresponding values of *N*_th_ for average distances between thermals of 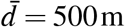, 1750 m, 3000 m were *N*_th_ = 40000, 3265, 1111, respectively.

We subsequently ran 500 simulations for each unique set of parameters, each in a randomly generated patchy environment (according to thermal distribution, activity, and occupancy), while the initial position and height of the bird and the goal location remained fixed. It is under these conditions that we ran our sensitivity analyses to quantify how each parameter affected the model outcomes, namely (1) success rate and (2) movement cost, described in the next section.

#### Focal simulations – static social environment

We then investigated the parameters *s, r* and *p* in more detail at 0.1 increments, with *p* ∈ [0.6, 0.7, 0.8, 0.9, 1.0] as lower predictability values were excluded following the results from the sensitivity analysis, where agents were unable to reliably reach the goal. We ran 2000 simulations for each unique set of parameters to explore the combined effects of those three parameters.

#### Collective simulations – dynamic social environment

We ran simulations for a homogeneous population of 100 individuals, with a shared overall goal location. Here we explored three parameters: *s, r*, and *p*, keeping the number of thermals fixed at 3265. The birds started from a well-defined initial position on a grid of edge size 10 km situated in the bottom left corner of the 100 km *×* 100 km arena. For each unique set of parameters, we ran 300 simulations and quantified the metrics below for all 100 individuals across simulations, resulting in 30,000 data points.

### 2.3 Performance and energy metrics

For each simulation run, we recorded success and flight cost to evaluate each bird’s movement performance. Success is a binary value indicating whether the bird successfully reached the overall goal (success = 1) or failed by landing before arrival (success = 0). We used these values to compute the success rate as the percentage of successful runs for a specific set of parameters. The flight cost was measured as the time (in seconds) taken by the bird to soar and glide from the starting location to the goal without landing. In unsuccessful runs, where the bird reached the ground before the goal, the flight time does not represent a comparable performance metric, therefore we only report the average cost of successful runs.

After running all the simulations, we proceeded in calculating the energy expenditure for all runs, whether successful or not, in terms of Basal Metabolic Rate (BMR), assuming that flapping flight could be adopted to reach the goal in unsuccessful runs where the bird reached the ground. This energy metric calculates the energy used during flight, considering that soaring and gliding consume energy at a rate of 2 times BMR, while flapping flight consumes energy at 20 times BMR [25, 28, 36]. In our simulations, flapping was not directly modelled, however we calculated the overall energy expenditure for unsuccessful individuals considering both the amount of flapping required at each grounding event to flap to the next thermal and soar back to an altitude equal to the starting one, and the average amount of times the bird would land to the ground before reaching the overall goal. By incorporating these calculations, we obtained a comprehensive energetic metric that accounted for both successful and unsuccessful runs, providing a more accurate assessment of the flight performance for a given set of parameters.

First, we computed 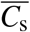, the average cost of successful individuals for each unique set of parameters. Next, for each single unsuccessful individual, we calculated 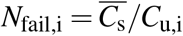, where *C*_u,i_ was the cost of the unsuccessful individual up to the failing point. This ratio estimated the number of failures that would occur over the time span of a successful run. We then evaluated the total energy expenditure for each bird *i* as:

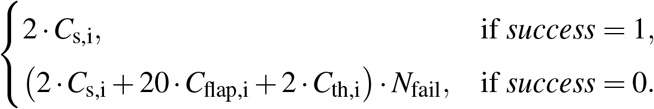

Here, *C*_flap,i_ represented the time the bird would have spent flapping between thermals if it had continued flying to reach the next thermal, and *C*_th,i_ the time spent in the next thermal in order to get back to the initial height.

After the initial sensitivity analysis, we fixed the number of thermals at *N*_th_ = 3265, therefore the average distance between thermals (Equation (5)) was 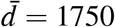 m. Using data from [37], we set the following values for the bird’s speed while flapping. The airspeed was *v*_f_ = 11 m/s, and the vertical ascent speed was *v*_fz_ = 0.7 m/s. We calculated the horizontal component of the velocity when flapping:

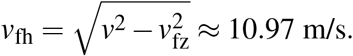

Next, we determined the time required to cover the distance to the next thermal:

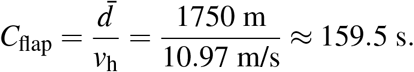

During this period of flapping, the bird gained altitude:

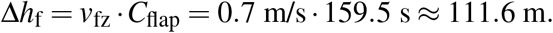

We then assumed that the bird would hit the next thermal with quality *q* = 2.5 m/s (average quality of active thermals) and would go back to the initial height of 1000 m in

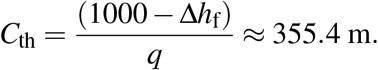

Finally, the “penalty” energy expenditure for each *N*_fail_ event resulted to be:

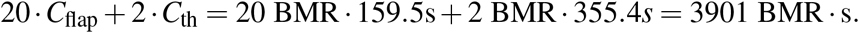

The energetic value of social information use was assessed by comparing the energy expenditure between a no-social baseline (*s* = 0) and social conditions (*s >* 0) under matched (*r, p, N*_th_) settings, for both static and dynamic social contexts.

## 3. Results

### 3.1 Sensitivity analysis

Systematic exploration of the parameters *s, r, p, a*_si_, and *N*_th_, while keeping others fixed, showed distinct patterns in the success rate (Fig. 3) which increases consistently with *s*, reaching a maximum at *s* = 1. In contrast, increasing *r* led to a decrease in success rate. For the predictability, higher values allowed for higher success rates, while low values (*p* = 0.4) resulted in a success rate smaller than 0.1. The availability of social information *a*_si_ had a weaker effect on the success rate, showing slight increases with a magnitude of change smaller than for the other parameters. When a high number of thermals (*N*_th_) were present, we observed a drastic drop, as very dense thermal fields reduce the efficiency of directed movement toward the goal: individuals tended to use many nearby thermals, leading to longer and less successful trajectories.

**Figure 3:**
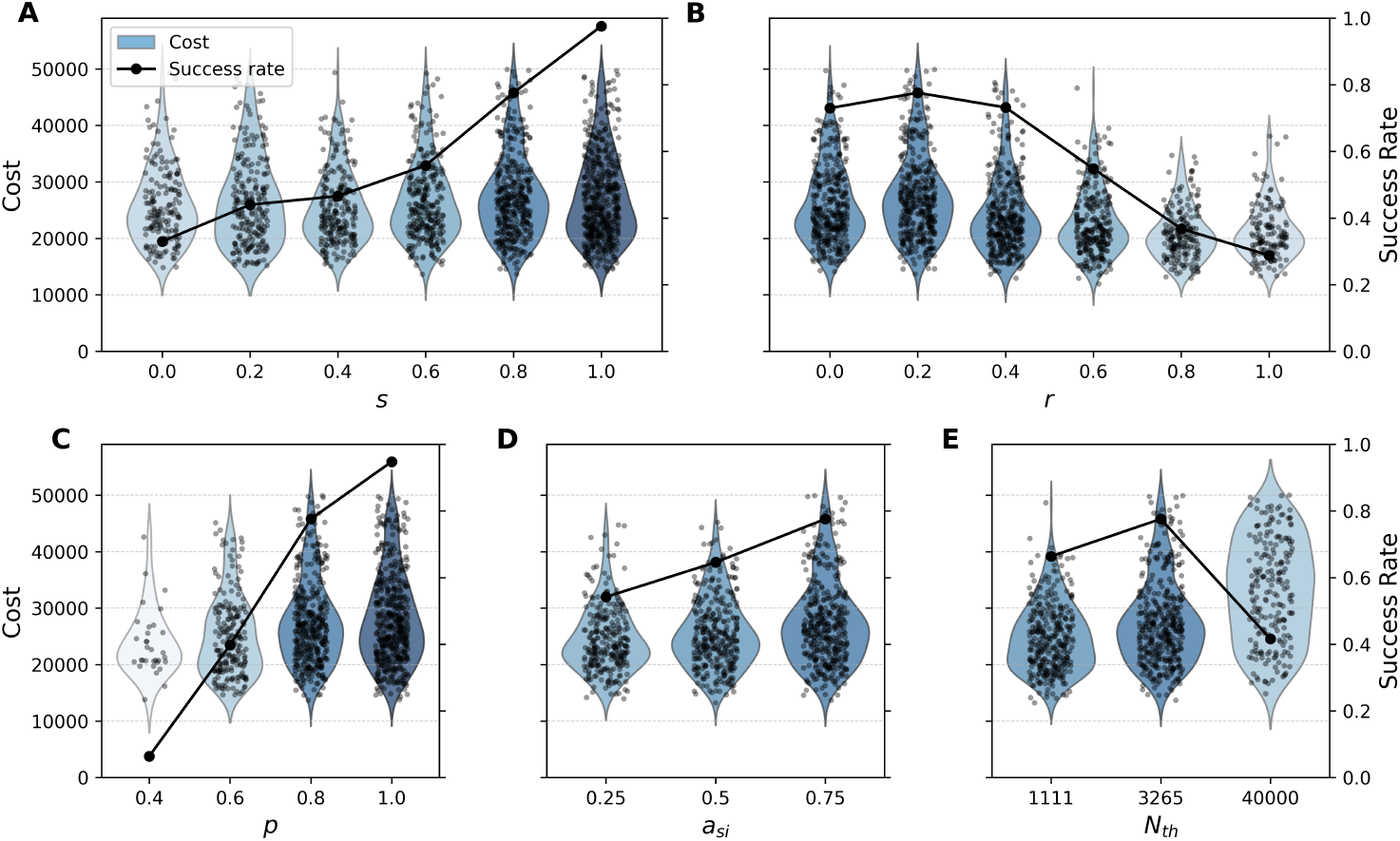
Sensitivity analysis showing the success rate and the cost distribution of successful runs as functions of five parameters: *s, r, p, a*_si_, and *N*_th_. In each panel, one parameter varies while the others are fixed: *s* = 0.8, *r* = 0.2, *p* = 0.8, *a*_si_ = 0.75, and *N*_th_ = 3265. Success rate increases with *s* (A), *p* (C), and *a*_si_ (D), but decreases with *r* (B). For *N*_th_ (E) it drops at very high values. The cost distribution is mostly stable, except at high *N*_th_, where the variance in cost increases. Colour indicates success rate (amount of points).

We found that predictability values below 0.6 were insufficient for achieving a high enough success rate to effectively analyse the simulation outcomes. For *p* = 0.4, the average success rate was below 0.5% for all the simulations with values of *s* smaller than 0.6. Hence, following the sensitivity analyses, we examined the simulation results on values of *p* between 0.6 and 1. The remaining environmental parameters were hereafter fixed at *a*_si_ = 0.5, as an intermediate level given its limited effect, and *N*_th_ = 3265, corresponding to a realistic average thermal density.

### 3.2 Focal simulations – static social environment

Overall, higher levels of risk can lower the average cost (Fig. 4), reducing the time required to reach the overall goal, but this benefit often comes at the expense of a reduced success rate across the simulations. As a consequence the average energetic cost to reach the goal increased with risk, as it takes into account the need to flap for a high number of unsuccessful simulation runs. In contrast, higher levels of sociality reduced energy consumption as it results in a greater number of successful simulation runs. Higher levels of environmental predictability reduce failure rates, but have only a limited effect on the average cost.

**Figure 4:**
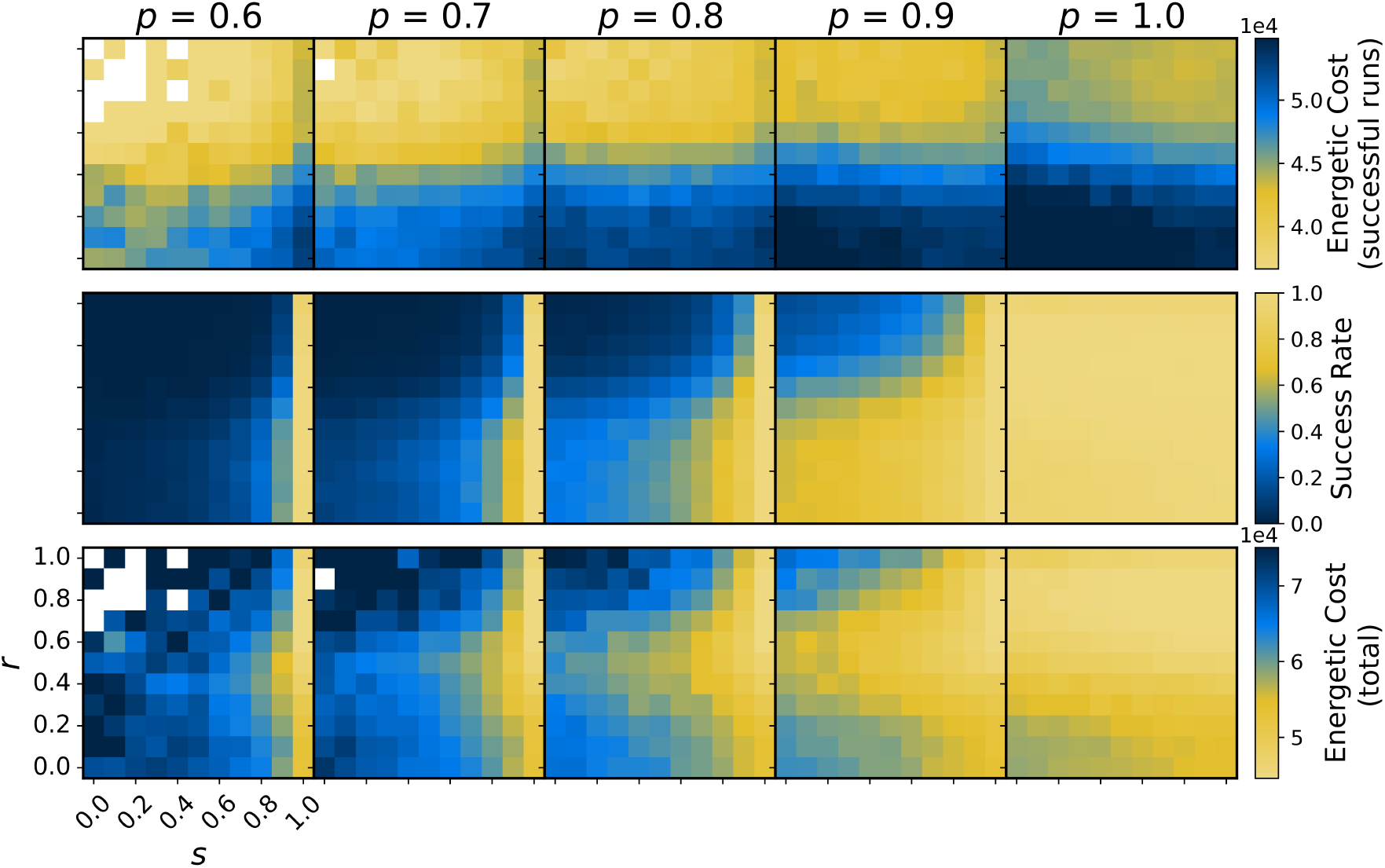
Average energetic cost of successful runs, success rate (percentage of successful runs), and energetic cost (with the inclusion of flapping flight) for different levels of predictability *p* (columns), sociality *s* (x-axis), and risk *r* (y-axis). Note that in the second row, the colour scale is inverted to match the logical interpretation of lighter colour as better performance. White cells indicate cases with 0% success rate, where no energetic cost could be calculated.

Performance in terms of energy consumption generally decreases monotonically with increasing *s* across all environmental conditions (Fig. 5), suggesting that the optimal choice is consistently *s* = 1. In contrast, energy consumption has a minimum at medium risk values. To quantify the energetic value of social information, we calculated the ratio between the energy consumption for a given value of *s* and the corresponding value when *s* = 0 (no use of social information) (Fig. 6). Relying on social information allows for reduced energy expenditure, especially in tough conditions when predictability is low (Table S1). For *p* = 0.7 and *r* = 0.3, having *s* = 1 allows for a 25.25% reduction in energetic cost compared to *s* = 0. For the same environmental condition, the percentage increases to 33.76% when *r* = 0.5 and to 36.67% for *r* = 0.7. When *p* = 1, all thermals are active and relying on social information may be expected useless, however, our simulations show that high values of sociality can nevertheless slightly reduce energy expenditure when travelling to a defined overall goal. While a very large number of available energy patch options can reduce movement efficiency by promoting less directed trajectories, the thermal density considered here does not reach such levels.

**Figure 5:**
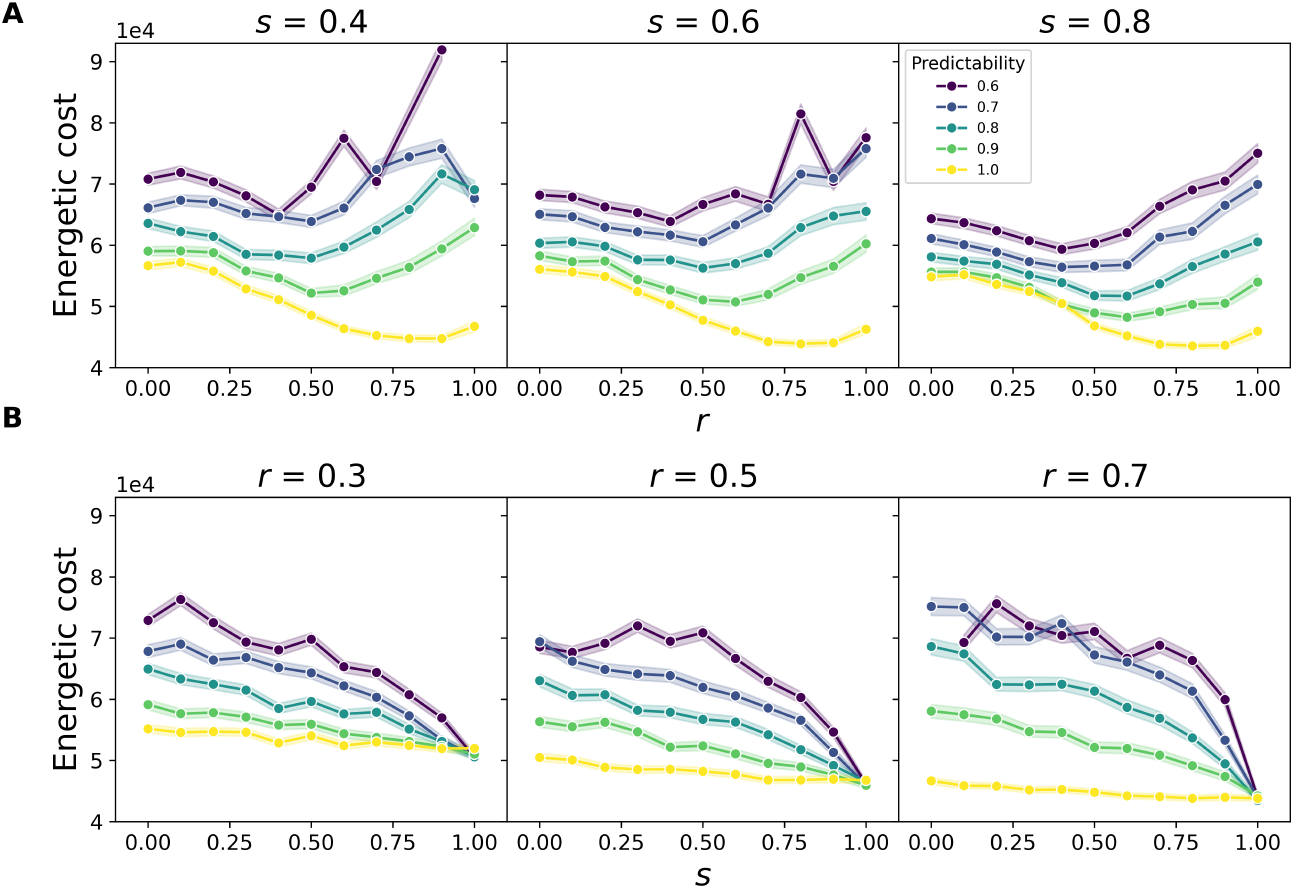
Trend of the energy cost over risk *r* (panel A) or sociality *s* (panel B) for different values of *s* or *r* (columns) and predictability conditions (line colours). Energetic cost reaches its minimum at intermediate *r* values and decreases monotonically with increasing *s*. This pattern is consistent across all predictability levels. The shaded area shows the 95% confidence interval. For low predictability, the large fluctuations in energetic cost result from an extremely low success rate on which energy expenses are calculated.

**Figure 6:**
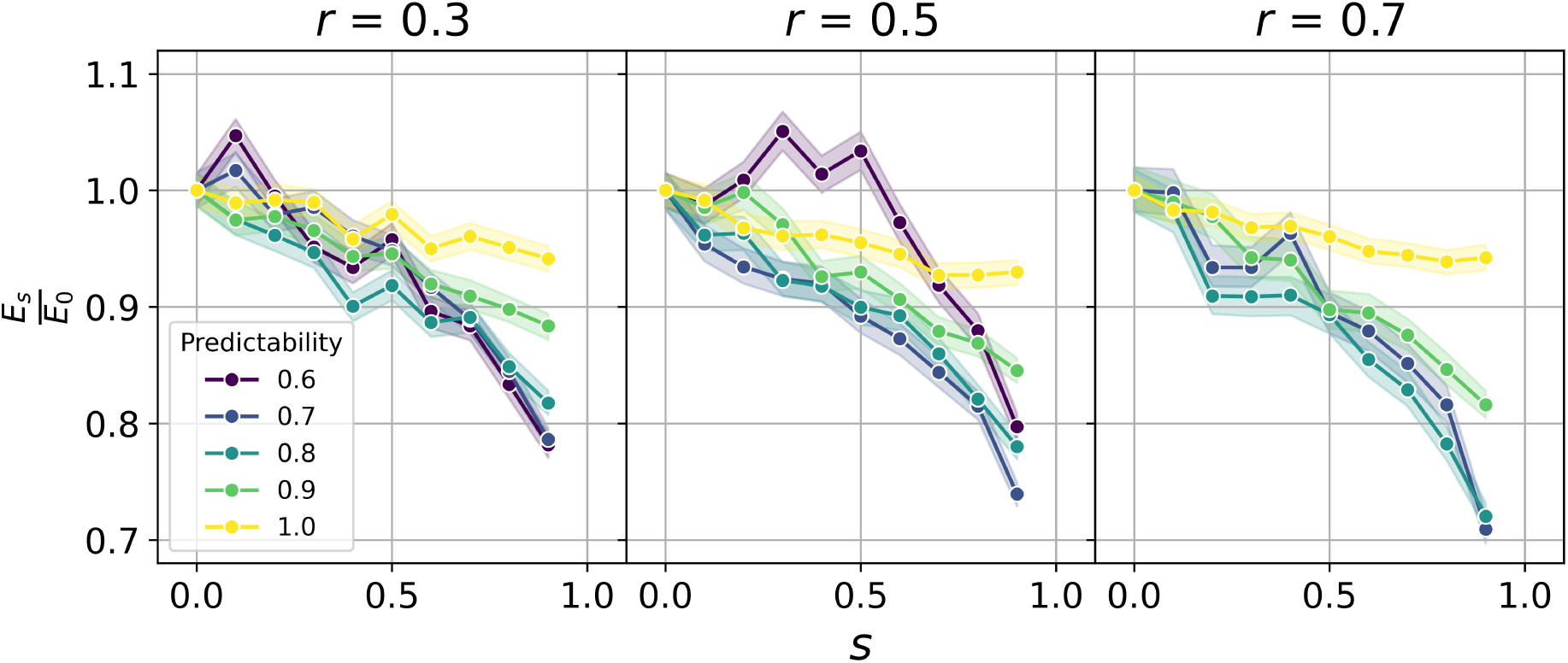
Ratio of the energy expenditure associated with a specific value of *s* (*E*_*s*_) over the value of energy expenditure in absence of sociality (*E*_0_) vs *s* for different values of *r* (columns) and predictability conditions (lines). The ratio has a general decreasing trend and the curves are steeper for lower values of *p*. At low predictability and low sociality, the patterns become more variable. The shaded area shows the 95% confidence interval.

### 3.3 Collective simulations – dynamic social environment

Building on the focal simulations, where a purely social strategy (*s* = 1) appeared optimal, we next considered the collective case to test whether this result holds in more realistic conditions. The collective model (Fig. 7) shows similar patterns to the static simulations regarding the effects of risk and predictability, and sociality generally reduces energetic cost. However, in contrast to the focal case, a fully social strategy (*s* = 1) leads to a marked increase in energetic cost. This demonstrates that while social information is usually beneficial, over-reliance on it can become suboptimal when all individuals rely exclusively on it (Table S2).

**Figure 7:**
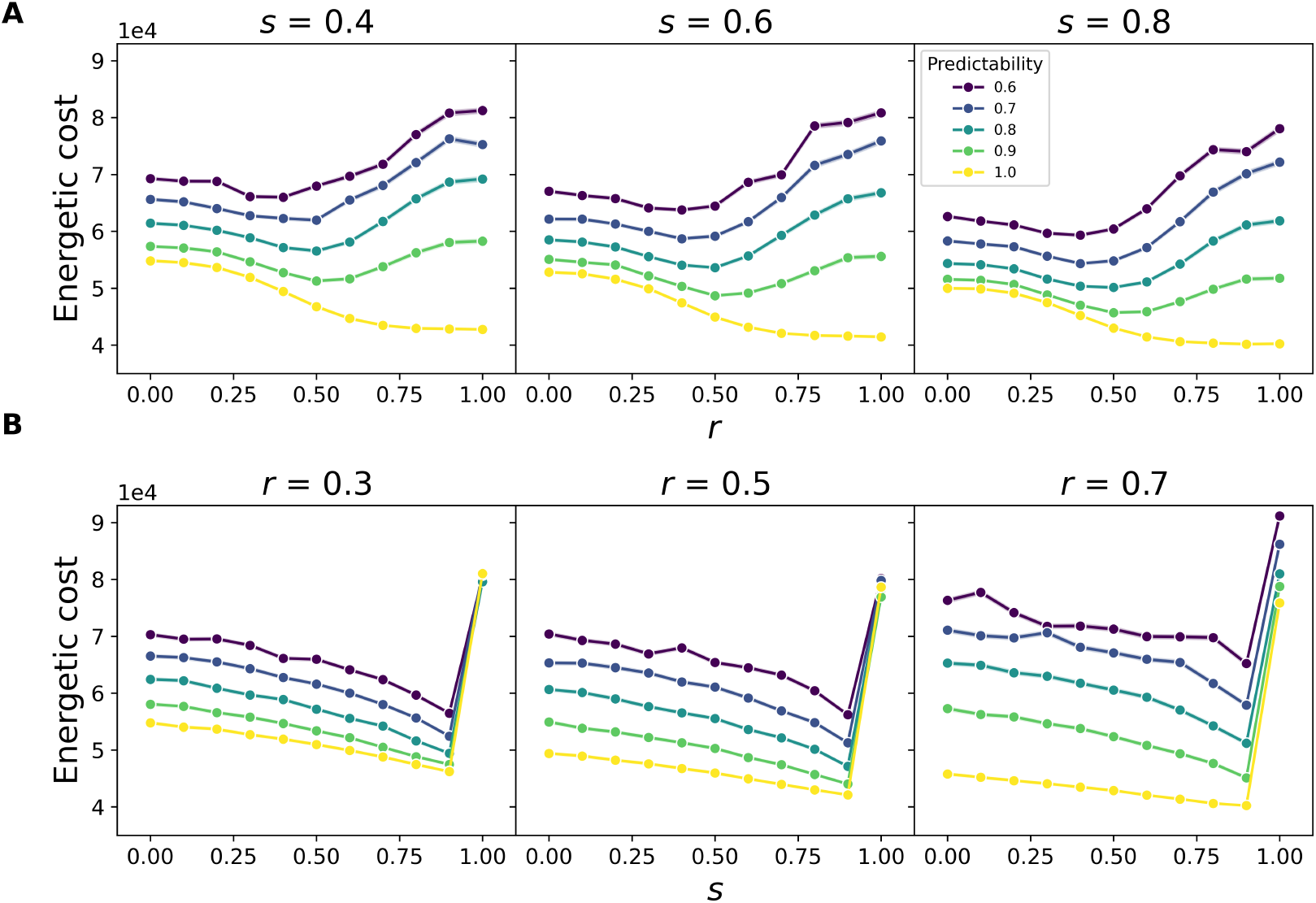
Trends of energetic cost over *r* (panel A) and *s* (panel B) for different values of *s* or *r* (columns) and predictability conditions (lines) in the collective model. In panel A, energetic cost decreases with increasing risk up to an inflection point just below *r* = 0.5, after which costs rise again, except when *p* = 1, where the minimum is reached at *r* = 1. In panel B, energetic cost generally decreases with increasing sociality, but shows a sharp increase at the extreme value *s* = 1. The shaded area shows the 95% confidence interval.

**Figure 8:**
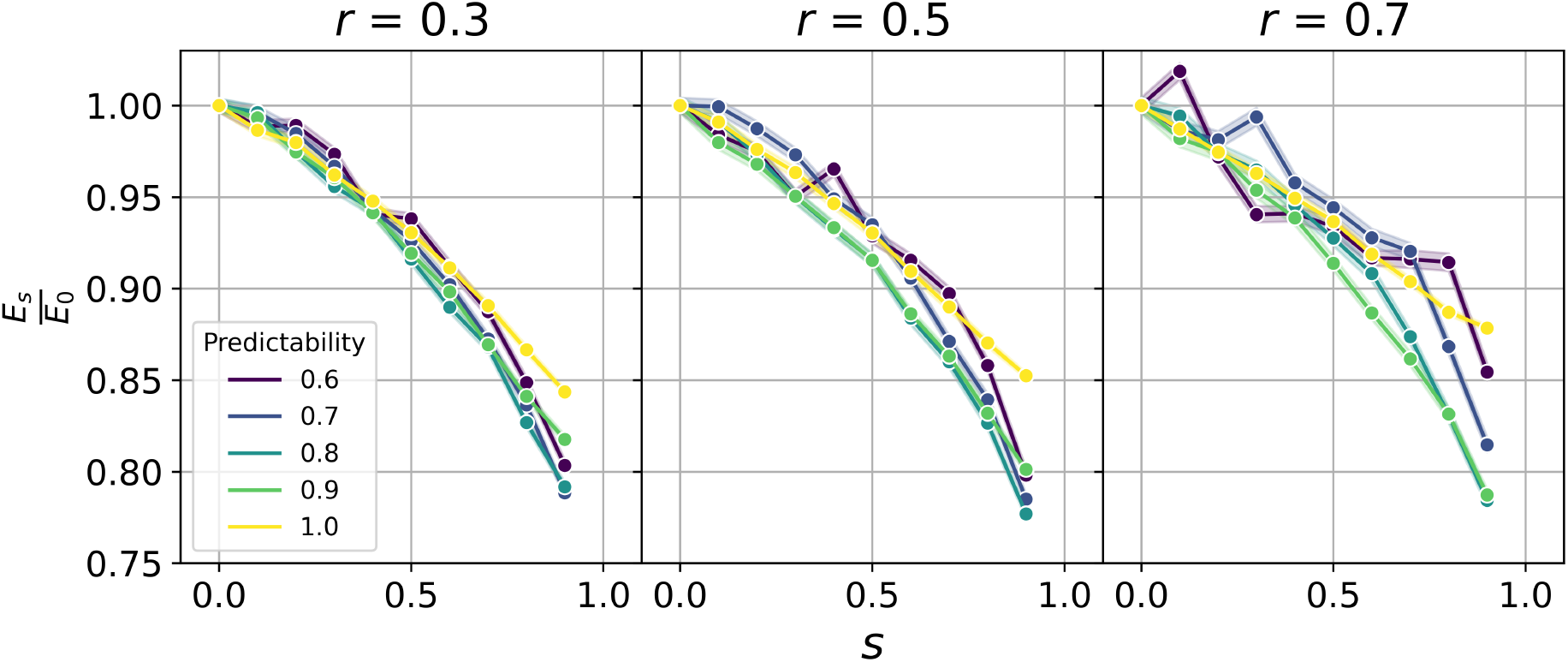
Ratio of the energy expenditure associated with a specific level of sociality *s* (*E*_*s*_) over the value of energy expenditure in absence of sociality (*E*_0_) vs *s* for different values of *r* (columns) and predictability conditions (lines) for the simulations of the collective model. The case of *s* = 1 is not represented in this plot. The shaded area shows the 95% confidence interval.

Compared to the focal case, the reduction in energy expenditure with increasing sociality is more pronounced, with a tighter relationship between *s* and *E*_*s*_*/E*_0_. The energetic value of social information, expressed as the ratio *E*_*s*_*/E*_0_, shows marked reductions with increasing *s*. For instance, at *r* = 0.3 and *p* = 0.7, increasing sociality from *s* = 0 to *s* = 0.9 reduces energetic cost by 21.15%. At *r* = 0.5, the reduction is 21.50%, while at *r* = 0.7 it lowers still to 18.54% (Table S2).

## 4. Discussion

Our computational model allows for the study of movement decisions in realistic physical environments. We consider soaring movement, characterised by a sequence of climbs in thermal updraughts and glides between them. While the classical speed-to-fly model (MacCready theory) provides an analytic optimum glide speed in order to maximise cross-country speed— assuming knowledge of the climb rate of the subsequent source of lift (thermal updraught) and the possibility to reach it [38]—in the real world, given the fine-scale dynamics of airflows [39], the environment is inherently uncertain. Thus, making decisions that rely on aeronautical rules alone can lead to poor choices and could involve extreme energetic costs, like the ones associated with flapping flight [25], as the strict assumptions of the theory may or may not be met. In our model we both create realistic environmental uncertainty and introduce social information in the decision-making process to cope with this uncertainty. We explicitly model height as a constraint affecting both kinematics and effective perception, making movement inherently three-dimensional; as a result, next-step decisions can be restricted to locations that are reachable solely by soaring, given the current altitude. Our model also includes a simple risk term that defines how height can be traded for speed. Together, these elements integrate physical realism, social information, and different risk strategies into a unified framework, linking aeronautical theory to social-information research, which otherwise, typically omits movement constraints [33, 7, 6, 40, 41].

In a dynamic world, an animal’s ability to move between resource patches depends not only on how it uses social cues from nearby individuals but also on the spatial distribution of those resources. We incorporate this aspect into our model by defining predictability as the probability that a thermal is active, reflecting the temporal variation in availability observed in nature. Although thermals exhibit some spatial predictability due to factors such as elevation, substrate type, or wind direction [42, 43, 44], whether a given thermal is active when a bird is within glide range remains highly variable. In our experimental setup, variation in the predictability parameter captures seasonal differences in thermal availability: thermals do not change state dynamically within a simulation; rather, thermal activity varies only across simulations, at a landscape level. We found that movement across the landscape becomes possible only when predictability is sufficiently high. In our realistically parametrised physical world, this threshold was identified at *p ∼* 0.6, corresponding to active thermals being, on average, *∼* 3.5 km apart. Greater average distances made it impossible for individuals to reach the goal under glide-polar constraints. At higher predictability levels, increasing the availability of social information (i.e. the probability that an active thermal is occupied by another individual) or thermal density had a limited effect. These results suggest that the scale over which individuals acquire and integrate information should therefore reflect the scale of predictability in patch availability and distribution.

We assume that social information reduces ambiguity [3] and operationalise its value as the reduction in energetic cost when individuals make movement decisions informed by the locations and states of others. Although we do not quantify uncertainty in mathematical terms, our computational model quantifies the energetic value of social information for movement in a patchy landscape and finds that relying on social information can yield energy savings up to 41% compared to not using it. For soaring-gliding flight, the energetic value of social cues is particularly pronounced given (i) the dynamic, invisible, and variable nature of airflows, (ii) the high energetic cost of misjudging a movement decision (i.e. errors trigger costly flapping flight), and (iii) the distinct movement signatures of gliding and soaring flight that present clear visual cues of the different underlying airflows. Therefore, the presence of social cues and the ability to acquire this information visually have the potential for a strong impact on a bird’s ability to cope with environmental uncertainty. In our model, movement decisions are probabilistic: acquired personal knowledge sets spatial priors, and social information updates beliefs about the current environmental state within the effective radius, altering the probability of selecting each location [22, 21]. We show that when social information is static and homogeneously available, movement costs decrease steadily with increasing sociality, and maximal sociality always yields the lowest costs across all levels of environmental predictability. Risk influences movement costs non-linearly, with intermediate risk generally minimising energetic expenditure, except when predictability is maximal, in which case high risk becomes the most efficient strategy. However, in scenarios of collective motion—where social cues indicative of resource availability arise dynamically—the advantage of maximal sociality disappears. Instead, the lowest energetic costs are achieved by combining high, but not extreme, sociality with intermediate risk.

At extreme levels of sociality in the collective scenario, individuals form tight clusters and their progress towards the goal slows down; consequently, overall efficiency decreases even though flapping remains rare. In real-world scenarios, regions with higher resource predictability, such as mountain ridges, are likely to concentrate both environmental resources and social information in specific areas. When social information is unevenly distributed or misaligned with the direction of an individual’s goal, relying exclusively on it may become disadvantageous. In such cases, social information might “pull” individuals away from their optimal path or intended goal, reducing movement efficiency, as observed in our collective scenario. This outcome suggests an explore-exploit trade-off, where remaining in the group reduces the probability of flapping, but limits the ability to exploit resources that would move an individual closer to the intended overall goal (e.g. a food resource distributed on a different landscape scale [33]). Our findings are consistent with trade-offs described in other collective systems, in which individuals must balance personally acquired environmental information with socially acquired information; studies of these systems show that excessive reliance on social cues can become suboptimal because it reduces independent sampling of the environment and increases the risk of propagating outdated or inaccurate information [45, 46]. Similarly, in ant foraging, strong reliance on collectively-built pheromone trails can limit exploration and lead to suboptimal path selection [47]. Introducing an explicit exploration parameter at the individual level could mitigate the counterproductive case of extreme sociality by avoiding excessive clustering of individuals, which would otherwise concentrate social information spatially. Moreover, extending the model to include variability in individual influence and responsiveness to others [48], would allow us to assess how different forms of heterogeneity among individuals affect movement efficiency.

Aligning information needs (i.e. uncertainty reduction) with environmental predictability enables energetically efficient movement strategies. However, long-term access to relevant social information typically requires remaining with the group, which may itself carry energetic costs. For example, relying on social information from conspecifics may increase competition for the resources they reveal. Our model assumes that thermals are non-depletable resources, meaning we did not account for potential competition within thermals (e.g., limited space within the thermal core where lift is strongest). Additionally, when movement is considered in the context of finding food patches or other depletable resources, competition extends beyond access to thermal updraughts and encompasses the target resource. In general, in groups, competition can become a critical factor. The benefit of following others therefore depends on a balance between energetic savings from improved movement and the costs of intensified competition for resources [49]. This balance between energy gained from resources and energy conserved during movement aligns with the principles of Optimal Movement Theory (OMT) [11], where efficient movement strategies are shaped by minimising energy consumption across different resource landscapes, and using social information to do so. Our results provide a baseline in the environmental energy landscape, showing that even with stable, non-depleting resources, the collective case does not favour full sociality. These findings suggest that under realistic conditions of resource depletion and competition, the optimal level of sociality would lie even further from the maximum. Similar findings were reported by [33], though without explicitly varying sociality, highlighting that the value of social information depends not only on movement efficiency but also on how competition for different resource types interacts with environmental conditions. In extreme environments with low resource availability and predictability, where space use may be limited [50, 51, 52], the payoff of social information for movement decision-making becomes more complex.

## 5. Conclusions

This study demonstrates the value of incorporating movement costs into an agent-based model of decision-making in dynamic landscapes explicitly. By linking energetic expenditure with information use, we show that social information can substantially reduce the costs of movement— but that its value depends critically on the distribution and predictability of resources. In patchy and unpredictable environments, the question becomes not only how much information is required, but whether the best strategy is to remain socially flexible, adjusting reliance on conspecific cues to match environmental dynamism. This raises the broader biological question of whether social information is a necessary adaptation for efficient movement in such landscapes, or whether its benefits are constrained when environmental structure imposes strong limits on movement opportunities.

Our work provides a flexible alternative to classical aeronautical models such as MacCready Theory, which prescribes an optimal glide speed between thermals but does so under strict assumptions and without accounting for socially informed decisions. By explicitly incorporating social information, risk-taking behaviour, and physical movement constraints, our approach moves beyond these limitations and offers new insights into how individuals and groups achieve efficient movement in patchy and unpredictable environments.

## Data accessibility

The code and the simulated data are archived on Zenodo [53]. A working version of the implementation code and further documentation is also available on [GitHub](https://github.com/eleonoragatti44/PatchyLandscapeMovement).

## Authors’ contributions

E.G.: Conceptualisation, Formal analysis, Methodology, Software, Validation, Visualisation, Writing – original draft, Writing – review & editing. H.J.W.: Conceptualisation, Funding acquisition, Project administration, Resources, Supervision, Methodology, Validation, Writing – review & editing. A.R.: Conceptualisation, Methodology, Validation, Writing – review & editing.

## Acknowledgments

This work was funded by VolkswagenStiftung Freigeist Fellowship (AZ-9B258 to H.J.W.) and E.G. is in part supported by IMPRS-QBEE. A.R. acknowledges support from the German DFG under Germany’s Excellence Strategy – EXC 2117–422037984.

