## Supplementary Tables for "Energetic benefits of social information for movement in patchy landscapes"

Table S1: Percentage of energy reduction for various values of $p$, $r$, and $s$ in the focal scenario compared to $s = 0$.

| p | r | $s = 0.5$ (%) | $s = 1.0$ (%) |
|---|---|---|---|
| 0.6 | 0.3 | 4.22 | 30.61 |
| | 0.5 | -3.38 | 32.65 |
| | 0.7 | – | – |
| 0.7 | 0.3 | 5.16 | 25.25 |
| | 0.5 | 10.79 | 33.76 |
| | 0.7 | 10.51 | 41.50 |
| 0.8 | 0.3 | 8.16 | 22.11 |
| | 0.5 | 10.03 | 26.26 |
| | 0.7 | 10.67 | 36.67 |
| 0.9 | 0.3 | 5.42 | 13.71 |
| | 0.5 | 7.02 | 18.47 |
| | 0.7 | 10.22 | 23.82 |
| 1.0 | 0.3 | 2.06 | 5.78 |
| | 0.5 | 4.49 | 7.37 |
| | 0.7 | 3.96 | 6.16 |

Table S2: Percentage of energy reduction for various values of $p$, $r$, and $s$ in the collective scenario compared to $s = 0$.

| p | r | $s = 0.5$ (%) | $s = 0.9$ (%) | $s = 1.0$ (%) |
|---|---|---|---|---|
| 0.6 | 0.3 | 6.19 | 19.66 | -14.52 |
| | 0.5 | 7.13 | 20.19 | -13.82 |
| | 0.7 | 6.62 | 14.56 | -19.46 |
| 0.7 | 0.3 | 7.37 | 21.15 | -21.90 |
| | 0.5 | 6.51 | 21.50 | -22.20 |
| | 0.7 | 5.58 | 18.54 | -21.29 |
| 0.8 | 0.3 | 8.38 | 20.83 | -27.46 |
| | 0.5 | 8.42 | 22.30 | -28.99 |
| | 0.7 | 7.25 | 21.57 | -24.07 |
| 0.9 | 0.3 | 8.08 | 18.23 | -39.17 |
| | 0.5 | 8.46 | 19.87 | -40.00 |
| | 0.7 | 8.63 | 21.26 | -37.48 |
| 1.0 | 0.3 | 6.94 | 15.64 | -47.95 |
| | 0.5 | 6.96 | 14.75 | -59.19 |
| | 0.7 | 6.33 | 12.16 | -65.68 |
